# Formation of recurring transient Ca^2+^-based intercellular communities during *Drosophila* hematopoiesis

**DOI:** 10.1101/2023.11.25.568594

**Authors:** Saar Ben David, Kevin Y.L. Ho, Guy Tanentzapf, Assaf Zaritsky

## Abstract

Tissue development occurs through a complex interplay between many individual cells. Yet, the fundamental question of how collective tissue behavior emerges from heterogeneous and noisy information processing and transfer at the single-cell level remains unknown. Here, we reveal that tissue scale signaling regulation can arise from local gap-junction mediated cell-cell signaling through the spatiotemporal establishment of an intermediate-scale of transient multicellular communication communities over the course of tissue development. We demonstrated this intermediate scale of emergent signaling using Ca^2+^ signaling in the intact, ex vivo cultured, live developing *Drosophila* hematopoietic organ, the Lymph Gland (LG). Recurrent activation of these transient signaling communities defined self-organized signaling “hotspots” that receive and transmit information to facilitate repetitive interactions with non-hotspot neighbors, transfer information across cells, and regulate the developmental progression of hotspots. Overall, this work bridges the scales between single-cell and emergent group behavior providing key mechanistic insight into how cells establish tissue-scale communication networks.

**Significance statement:** Cells coordinate their internal state and behavior by exchanging information with other cells in their vicinity. These local interactions are integrated across space and time to enable synchronized function at the tissue scale. Using live microscopy imaging of the *Drosophila* Lymph Gland, and by applying computational analyses, we identified and characterized a new mode of cellular communication through self-organized recurring coordinated short-term activation at the intermediate scale of 3-8 cells, which we call “hotspots”. We reveal that hotspots form over the course of tissue development, and are dependent on specific proteins, called gap-junctions, that enable communication between adjacent cells. Hotspots repeatedly transmit and retrieve information to and from their non-hotspot neighbors to spread information throughout the tissue to regulate and coordinate tissue function.

## Introduction

The emergence of collective cell behavior is an essential component of many basic biological phenomena such as tissue morphogenesis (1), cell migration (2), or bacterial quorum sensing (3, 4). Key to understanding collective cell decision-making is elucidating how local information transfer between cells is integrated in space and time. This spatial and temporal integration of information is essential for regulating the emergence of collective behavior at the multicellular scale (5, 6). The *Drosophila* hematopoietic organ, the Lymph Gland (LG), is a powerful, genetically tractable, model to study how information is integrated in space and time to facilitate collective cell behavior. The LG contains dozens of stem cell-like blood progenitors that are largely quiescent but can be collectively activated in certain conditions, such as in response to pathogenic infection, to rapidly produce hundreds of highly differentiated blood cells with infection-fighting characteristics (7, 8). Long-term culture and live imaging of the intact LG showed that Calcium (Ca^2+^) signaling, which is transmitted between blood progenitor cells through gap-junctions, mediated essential information transfer across large distances in the LG (9). Ca^2+^ levels serve a key function in controlling blood progenitor fate as the activity of multiple pathways that regulate progenitor behavior, including JAK/STAT and CaMKII signaling, is modulated by the amount of Ca^2+^ in the cell at a specific time (9, 10). Gap junctions, intracellular channels that directly link adjacent cells to allow them to exchange ions and other small molecules, can help cells form signaling networks (9, 11, 12). In characterizing, at the population scale, the gap-junctions based, Ca^2+^-mediated, multicellular signaling network in the LG we observed synchronized cell pairs that were located up to 38 cell diameters (∼190 µm) from one another. Importantly, functional studies illustrated that the gap-junction mediated Ca^2+^-signaling network was required for proper regulation and function of the LG by coordinating fate decisions at the population scale (9). A key question that emerged from our previous results was how the local information transfer between adjacent cell pairs formed a global multicellular network. Specifically, we wanted to characterize and understand the intermediate stages that allowed cell-cell signaling exchanged between individual cells to become collective signaling.

Here we identified, using spatiotemporal analysis of Ca^2+^-signaling in live intact LGs, the gradual formation of communicating communities of 3-14 progenitor cells over the course of development, through intercellular gap junction-mediated signaling. Recurrent signaling activity of these communities formed hotspots of local information transmission highlighting heterogeneity in intercellular information transfer as a potential contributor to collective decision making. Taken together, our results explain how the exchange of information between individual cells in the *Drosophila* LG becomes an emergent behavior involving multiple cells. This provides insight into the bridging of the scales between single-cell and emergent group behavior.

## Results

### Propagating intercellular Ca^2+^ signaling forms communicating communities in the *Drosophila* lymph gland

We investigated Ca^2+^ signaling in individual blood progenitors using live imaging of intact, *ex vivo* cultured, LGs (Fig. 1A). By manual qualitative selection of adjacent blood progenitor pairs, we previously showed that Ca^2+^ signals propagate between neighboring blood progenitor pairs and this propagation is mediated by gap junctions (Video S1) (9). To systematically and quantitatively characterize the patterns of signal synchronization across scales in-depth, we measured the temporal correlation between Ca^2+^ signals in all blood progenitor pairs in the LG. This analysis identified a negative correlation between the distance between blood progenitor pairs (termed *cell pair distance*) and the level of coordination in their Ca^2+^ signals (termed *cell pair correlation*). This means that, on average, closer blood progenitor pairs were more synchronized in terms of Ca^2+^ signaling than distant pairs (Fig. 1B, Fig. S1A). These data identified a sub-population of highly synchronized cell pairs, where cells were located within a distance of approximately 14 µm from one another, about two cell diameters apart. Indeed, partitioning the data to close (≤ 14 µm) versus far (≥ 14 µm) cell pairs showed that close pairs were more likely to be in a higher level of synchronization (Fig. 1C, Fig. S1B). This subpopulation of highly synchronized close-cell pairs highlighted the heterogeneity in cell-cell information transfer. However, it was still unclear how this local cell-cell synchronization propagates from the scale of cell pairs to the multicellular scale.

**Figure 1.**
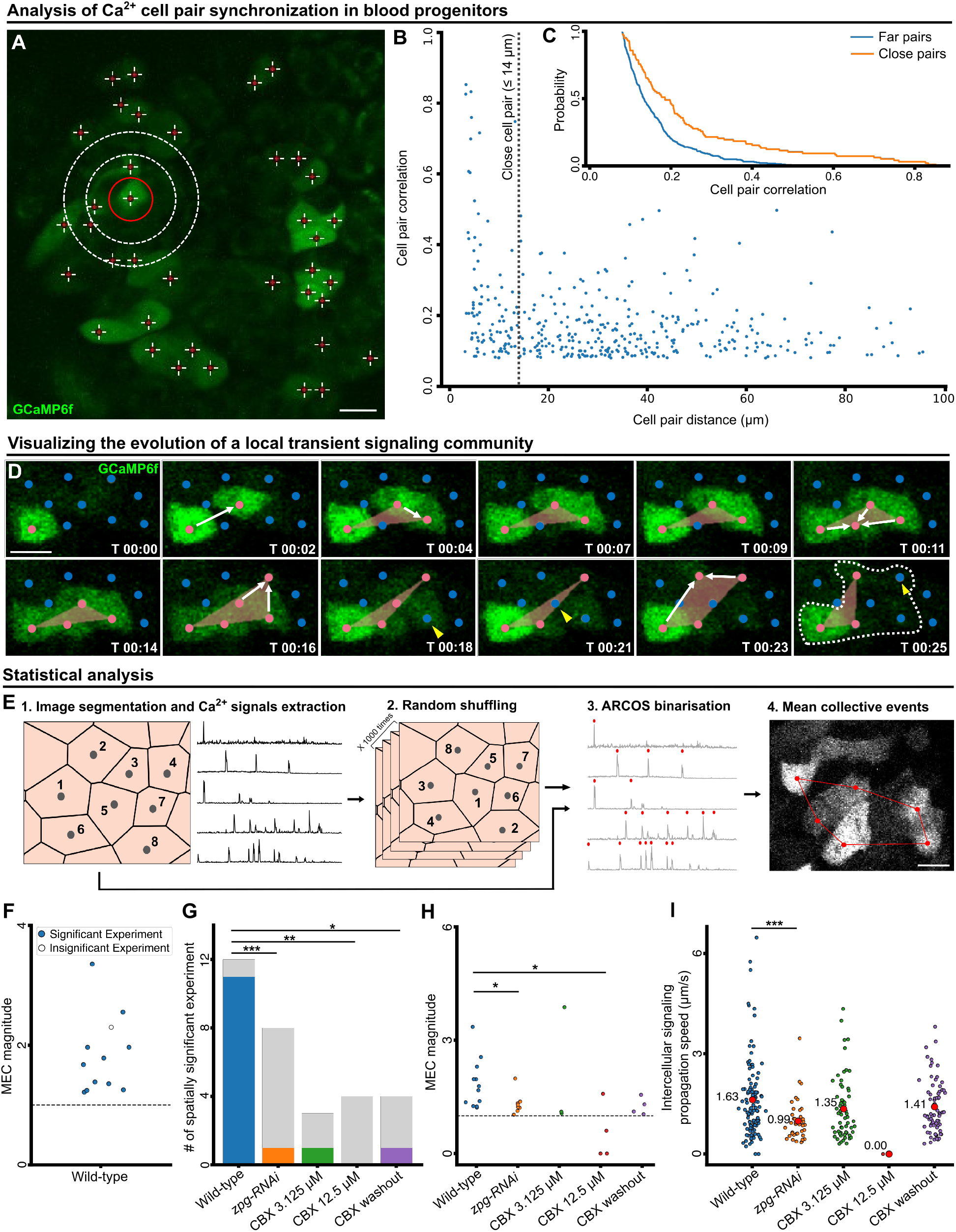
Blood progenitor cell-cell communication forms communities of propagative Ca^2+^ signaling. **(A)** Representative confocal image showing Ca^2+^ signaling activities in blood progenitors of a LG visualized using GCaMP6f (in green). Red crosses indicate the center of individual cells. White circles indicate adjacent blood progenitors to the cell marked by the red circle, at distances of 7 µm and 14 µm from it correspondingly (see also Video S1). Scale bar = 10 µm. **(B)** Spatial analysis of blood progenitor pairs that showed a statistically significant correlation (p < 0.05) in their temporal Ca^2+^ signals. Each data point (blue) represents a cell pair. Cell pairs Ca^2+^ Pearson correlation was correlated with the cell pairs distance. *N_cells_* = 57, *N_pairs_* = 385, Pearson correlation between cell pair Ca^2+^ correlation and distance = −0.244, p-value = 0.003. See also Fig. S1A for an analysis of all cell pairs. **(C)** Cumulative distribution of Pearson correlation of the close (orange; N = 98, µ = 0.246, σ = 0.187) and far (blue; N = 287, µ = 0.160, σ = 0.084), significantly Ca^2+^ correlated blood progenitor pairs (same pairs as in B). Each value *F_g_*(*x*) in the plot is the probability of a pair in group *g* to have a Pearson correlation coefficient greater than *x*. Kruskal-Wallis statistical test verified a significant difference between the two distributions (p-value < 0.0001). See also Fig. S1B for an analysis of all cell pairs. **(D)** Representative confocal images showing a Ca^2+^ signaling propagation event, detected by ARCOS, which defined a transient community involving 6 blood progenitors (see Results text and Methods). GCaMP6f is labeled in green. The center of each cell is marked in red (active, i.e., showing Ca^2+^influx) or blue (inactive). Time (T, in second) is annotated in each frame. Orange polygons visualize the cell centers transiently participating in a community in each frame. White arrows indicate the inclusion of new activated cells in the community, yellow arrowheads indicate the deactivation and exclusion of cells from the community. All the cells that participate in the community throughout its evolution are marked in the last frame (T = 00:25) in a dashed white polygon. Scale bar = 5 µm. **(E)** Schematic of the spatial shuffling analysis (see also Methods). (1) Single-cell segmentation and extraction of Ca^2+^time series. (2) Random spatial shuffling of the Ca^2+^ time series of all cells, repeated 1000 times, correspondingly generating spatially permuted experiments. (3) ARCOS binarization: Ca^2+^ peak detection (red). (4) ARCOS community detection (red, white is GCaMP6f). Recording of the mean collective events per cell (MEC) and statistical comparison of MEC for observed versus *in silico* permuted experiments. Scale bar = 5 µm. **(F)** Analysis of MEC magnitude (N = 12 LGs). Mean ratio between MEC of the observed and the *in silico* permuted experiments. The ratio of value 1 (dashed horizontal line) implies no change in the magnitude. The bootstrapping significance test showed spatial significance for 11/12 LGs (color-filled circles). **(G-I)** Gap junction inhibition experiments. Wild-type LGs (N = 12), RNAi-mediated *zpg* knockdown (N = 8), 3.125 µM CBX (N = 3), 12.5 µM CBX (N = 4), and CBX washout (N = 4). Statistical analyses: * - p < 0.05, ** - p < 0.01, *** - p < 0.001, **** - p < 0.0001. (**G**) Spatially significant experiments. For each experimental condition, gray indicates the number of insignificant and color indicates the number of significant LGs. Significance was determined using Fisher’s exact test. **(H)** Analysis of MEC magnitude. Each data point corresponds to one LG. Significance was determined using the Kruskal-Wallis test to evaluate the differences between the wild-type and the other conditions. **(I)** Analysis of intercellular signaling propagation speed between adjacent cells in a community. Each data point (red) represents the average cell-cell signaling propagation speed calculated according to the relative activation timing between adjacent pairs in each transient community (see Methods). Wild-type (N = 113 communities, mean information spread µ = 1.63 µm/second), RNAi-mediated *zpg* knockdown (N = 39, µ = 0.99 µm/second), 3.125 µM CBX (N = 62, µ = 1.35 µm/sec), 12.5 µM CBX (N = 1, µ = 0 µm/second), and CBX washout (N = 71, µ = 1.41 µm/second). Statistical significance was determined using the Kruskal-Wallis test to evaluate the differences between the wild-type and the other conditions.

To detect and quantify collective spatiotemporal signaling events, i.e., signaling events that involve more than two cells, we applied a computational method known as the “Automatic Recognition of COllective Signaling” (ARCOS) (13). ARCOS binarizes the single blood progenitor Ca^2+^ signal, according to its magnitude, to “active” (Ca^2+^ peak) or “inactive”, followed by spatiotemporal clustering of cells that are synchronously active (peaks ≤ 15 seconds apart) (13). This analysis defines “collective signaling events” that we refer to as local transient *communities* of blood progenitor signaling (Video S2). Every community consists of a minimum of three cells that were active simultaneously or within a 15-second delay. Using ARCOS, we were able to monitor the formation and disintegration of a community (Fig. 1D, Video S3): following an initial Ca^2+^ spike, subsequent activation of adjacent blood progenitors, as marked by red dots connected by a white arrow, initiated a 3-cell community (Fig. 1D, 0-7 seconds). The community gradually grew, which was observed as Ca^2+^ activation in adjacent cells (Fig. 1D, red dots and white arrows, 11-23 seconds) and shrunk by deactivation of cells in the community (Fig. 1D, yellow arrowheads, 18-25 seconds). Throughout its evolution, this community involved 6 cells (Fig. 1D, marked by a white dashed polygon, 25 seconds) with a maximum of 5 cells being active simultaneously (Fig. 1D, 16 seconds). Our analysis identified communities of local intercellular transfer of signaling information involving 3-14 blood progenitors per community, with a median community size of 4 cells and 30% of communities having at least 5 participating cells (Fig. S2, example in Video S3). Two potential confounders of this analysis were the stochastic co-incidence of activation events and the presence of areas with higher local cell densities, both of which may lead to the detection of spurious collective signaling events by ARCOS (Fig. S3). To mitigate these potentially confounding factors, we spatially shuffled the cells (i.e., randomized their location), applied ARCOS to identify collective signaling events in the spatially permuted experiment, and recorded the <u>m</u>ean number of collective signaling <u>e</u>vents per <u>c</u>ell (mean events per cell, MEC) across the entire population. We repeated the sequence of random shuffling and ARCOS analysis 1,000 times (Fig. 1E) and recorded: (A) the statistical significance - the fraction of times that the MEC of these *in silico* spatially permuted experiments were equal or exceeded the MEC of the observed (un-permuted) experiment, and (B) the magnitude - the mean ratio between the experimentally observed MEC and each of the *in silico* spatially permuted MEC. All replicates, but one (11/12), showed significant elevation in magnitude of MEC, by a factor of 1.2-3.3 fold in respect to the *in silico* permuted experiments, indicating that the collective signaling events were a local property of this multicellular system (Fig. 1F). Altogether, our data suggests that local cell-cell information transfer integrates in space and time to form multicellular communities of Ca^2+^ signal propagating blood progenitors in live intact *ex vivo* cultured LGs.

### Gap junctions mediate the propagation of Ca^2+^ signals in blood progenitor communities

We previously demonstrated that gap junctions were required for cell-to-cell Ca^2+^ propagation between the blood progenitors in the *Drosophila* LG (9). To assess the role of gap junctions in the formation of intercellular communities, we analyzed *ex vivo* cultured LGs using live imaging under different conditions where gap junctions were perturbed. Specifically, we used both a genetic and a pharmaceutical-based approach to disrupt gap junction-mediated communication between blood progenitors. First, we used an RNA interference (RNAi) approach to knock down the expression of the gap junction protein Innexin 4, known by its gene name *zero population growth,* or *zpg* (14). We have previously shown that Zpg is the main gap junction channel mediating Ca^2+^ signaling between blood progenitors (9). Second, we used the gap-junction blocker known as carbenoxolone (CBX). We performed RNAi-mediated knockdown of *zpg* (N = 8), a low dose CBX treatment (3.125 µM; N = 3), a high dose CBX treatment (12.5 µM, N = 4), or a control where we first treated with 100 µM CBX and then washed it out (N = 4). Analysis of these different treatment groups showed that gap-junction inhibition led to a drastic decrease in the fraction of experiments with significant local communities (Fig. 1G), the magnitude of collective signaling communities (Fig. 1H), and the intercellular signaling propagation speed between adjacent cells (Fig. 1I). Intriguingly, washout experiments that were previously shown to rescue the network properties and cell-cell propagation (9), did not rescue the fraction of collective signaling-event communities (Fig. 1G), but did rescue the magnitude of communities (Fig. 1H) and the intercellular signaling propagation speed between adjacent cells in a transient community (termed *intercellular signaling propagation speed*, Fig. 1I). This suggests that perturbation of gap junction-mediated communication may have a long-lasting effect on the signaling community that persists even after CBX is removed. The association between the LG’s mean cell activation rate (i.e., frequency of cell activation), mean local cell density, and the MEC rate, meaning the mean frequency that a cell participates in a transient community, were maintained for most gap junction inhibition perturbations (Fig. S4A-C). However, Zpg depletion (using RNAi) or inhibition (using CBX) led to increased cell activation, i.e., higher frequency of Ca^2+^ spikes, but reduced MEC rate for the same activation level (Fig. S4D), suggesting a compensation mechanism where Zpg-depleted or inhibited cells try to compensate for reduced cell-cell communication capacity by increasing their activity. These results validate the critical role of gap junctions in the formation of Ca^2+^-based intercellular communities.

### Recurrent activation of communication communities forms hotspots of local information processing hubs

We next asked whether the same cells participate in multiple (transient) signaling communities, which, if true, could suggest that these communities act as signaling communication “hubs” that repeatedly receive and spread information to synchronize the multicellular network. To quantitatively assess this possibility in wild-type LGs, we recorded for each cell the number of times it participated in signaling communities. Visualization of the number of times each cell participated in a community revealed spatial heterogeneity with recurrent activation of specific communities, that we call “*hotspots*”, involving groups of spatially adjacent cells with enriched participation in signaling communities with respect to the population (Fig. 2A, Fig. S5A-E, criteria for hotspot identification are detailed in the Methods). We identified hotspots that met these criteria in 9 out of the 12 wild-type (non-treated) LGs. The number of hotspots per LG ranged between 1 to 3 with each containing between 3 to 15 cells. To verify that hotspots were not a mere consequence of increased cell activation we devised a bootstrapping-based statistical test (Fig. 2B-D). First, we matched and replaced at least 50% of the cells in the hotspot with other cells in the same experiment that did not take part in the hotspot and had, at minimum, the same amount of activations. Second, we switched the Ca^2+^ time series for each pair of matched hotspot and non-hotspot cells, and then detected collective signaling events in this *in silico,* spatially permuted, experiment (Fig. 2D). Third, we recorded the MECs for cells participating in the hotspot of the *in silico* permuted experiment. We repeated these steps of switching “hotspot” with non-hotspot cells with at least the same number of Ca^2+^ activation, up to 1,000 times for each hotspot, recorded the difference between experimentally observed hotspots and their *in silico* permuted versions, and determined the statistical significance. Statistical significance was determined by calculating the fraction of permutations where the hotspot MEC values in the *in silico* experiments were equal to or exceeded the MEC values of the observed (wild-type, non-permuted) experiment. This analysis showed a dramatic decrease in the MEC following spatial permutation (Fig. S5), statistically validating 8 of 14 hotspots, spread over 5 of the 12 live intact *ex vivo* cultured LGs (Fig. 2E). Qualitative observation of the validated hotspots locations did not identify a typical spatial pattern in respect to the LGs. Gap-junction perturbations, even after washout, showed reduced numbers of validated hotspots (Fig. 2E).

**Figure 2.**
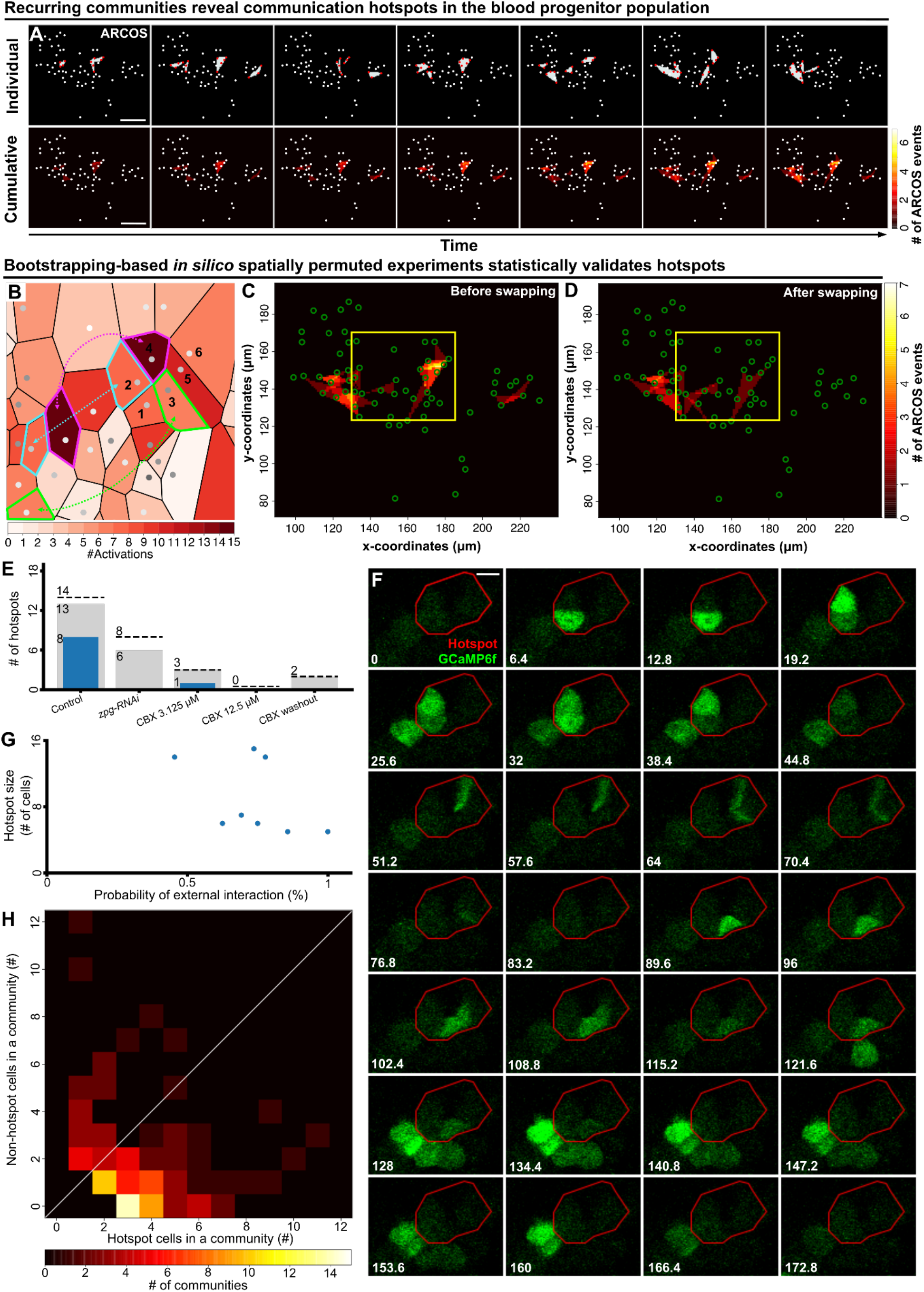
Recurrent activation of communities forms hotspots that act as local information hubs. **(A)** Representative time-lapse images showing the formation of hotspots over time. A hotspot is defined by recurring transient communities (see Methods). Top panels: transient communities (marked by colored polygons, red dots mark activated blood progenitors) in a wild-type LG. Bottom panels: the integrated number of transient communities over time. Each white dot represents an individual blood progenitor. Each panel corresponds to its matching top panel. Scale bar = 15 µm. **(B)** Single-cell Voronoi tessellation, corresponding to the yellow region of interest shown in panels C and D, and illustrating the bootstrapping-based *in silico* permutation experiment (see Methods). The color of each cell (polygon) reflects the number of activations (i.e., calcium spikes) each cell exhibits. Six cells that participate in a hotspot are numbered and dashed color-matched arrows indicate cell swapping. The swapping is performed for cell pairs with similar activation, where one cell is within and the other outside the hotspot (see Methods). **(C-D)** Representative field of view showing the integrated number of transient communities each cell participated in over time (#ARCOS events) before (C) and after (D) *in silico* permutation (see B). Green circles: the center location of each blood progenitor. Brighter areas indicate more occurrences of communities. The yellow region of interest marks the hotspot that is also shown in B. **(E)** Hotspot statistics. Hotspots were pooled across experiments according to the experimental condition. Dashed line - pooled number of hotspots. Gray - pooled number of hotspots with sufficient data for statistical analysis. Blue - number of statistically significant validated hotspots. Hotspot significance was determined according to 100-1000 different *in silico* permutation experiments with a bootstrapping significance threshold of 0.05. **(F)** Time-lapse evolution of a representative hotspot. The hotspot was defined according to the integrated number of transient communities per cell across the experiment (red polygon; see Methods). Transient communities involve cells within and outside the hotspot. GCaMP6f labeled in green. Scale bar = 5 µm. **(G)** The probability of hotspot cells interacting with cells outside the hotspot through common transient communities as a function of the hotspot’s size (i.e., the number of cells in the hotspot). The analysis included the 8 statistically verified hotspots pooled across all wild-type LGs. **(H)** Histogram of the number of hotspot cells (x-axis) and non-hotspot cells (y-axis) in communities that define the hotspots - each observation used for this histogram is defined by a community. White diagonal (*y* = *x*) indicates an equal proportion between hotspot to non-hotspot cells.

Our findings raised two important questions regarding the interaction of hotspots with their environment. First, do hotspots function as self-contained groups of cells, interacting predominantly within their enclosed local surroundings? Second, do hotspots initiate the spread of information, or are they more responsive to incoming non-hotspots external signals? Following the evolution of a transient community showed alternating interactions between cells inside and outside a hotspot (Fig. 2F). To systematically decipher the interactions between hotspots and their surrounding environment we analyzed the spatiotemporal communication patterns of all the validated hotspots that were pooled across the wild-type LGs (N = 8 hotspot). To quantify the interactions of hotspots with their surrounding cells, we calculated each hotspot its probability of engaging with cells outside the hotspot through common transient communities (Methods). The majority of hotspots (7 out of 8) interacted with non-hotspot cells in more than half of their transient communities, this interaction was independent of the size of a hotspot (Fig. 2G), and was dominated by communities that involved 2-4 cells within the hotspot and 1-2 cells external to the hotspot (Fig. 2H). Specifically, 70% of hotspot communities had at least one non-hotspot cell involved, and 76% of these communities involved more hotspot cells than non-hotspot cells (Fig. 2H). These interactions of a hotspot with its surrounding cells did not have a systematic direction, starting from hotspot cells outwards or initiating externally from adjacent non-hotspot cells (Fig. S6, Methods). Furthermore, we did not identify cells that repeatedly initiated a hotspot’s transient communities, suggesting stochasticity in hotspot initiation. These observations established the existence of gap-junction-mediated communication “hotspots”, where recurrent Ca^2+^ communities coalesce into larger communication hubs that repeatedly spread and retrieve information throughout the blood progenitors.

### Gradual formation of communication communities and their recurrent activation during LG development

In flies, hematopoiesis is subject to developmental regulation, with blood progenitors exhibiting distinct behaviors at different larval stages (15, 16). Specifically, cell proliferation and differentiation show distinct patterns at different points along the developmental timeline (Fig. 3A) (16). For example, the differentiation of mature blood cells starts around the mid- to late-second instar transition and peaks around the mid-second to mid-third instar larval stages (10, 15, 16). The level of mature blood cell differentiation gradually declines as the LG develops and becomes significantly attenuated upon entry into the mid-third instar stage (Fig. 3A) (10, 16). In contrast, cell proliferation in the blood progenitors peaks earlier, during the first- to second-instar stages, when the progenitor repertoire rapidly expands (15, 16). Shortly after the onset of differentiation, the rate of cell proliferation slows down but remains active until the mid-third instar stage (Fig. 3A) (17).

**Figure 3.**
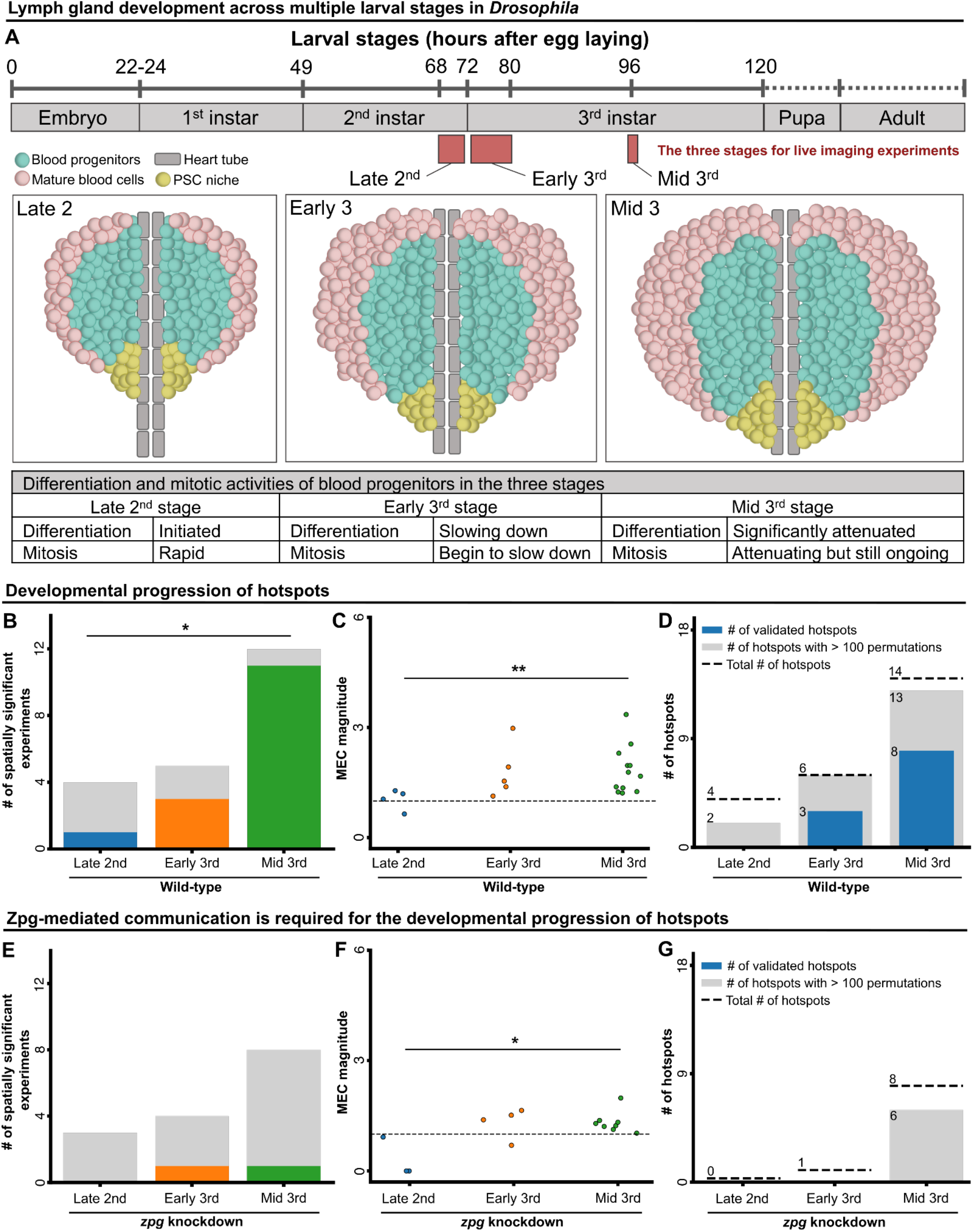
Gradual formation of communication communities during development. **(A)** Lymph gland development throughout *Drosophila* larval stages (see Methods). **(B-D)** Analyses of wild-type LGs from the late-second instar stage (N = 4), early-third instar stage (N = 5), and mid-third instar stage (N = 12). **(B)** Number of spatially significant experiments. For each experimental condition, gray indicates the number of insignificant and color indicates the number of significant LGs. Significance was determined using Fisher’s exact test. **(C)** MEC magnitude. Each data point corresponds to a single LG. Significance was determined using the Kruskal-Wallis test to evaluate the differences between the different developmental stages. **(D)** Hotspots statistics. Dashed line - pooled number of hotspots. Gray - pooled number of hotspots with sufficient data for statistical analysis. Blue - number of statistically significant validated hotspots. Hotspot significance was determined according to 100-1000 different *in silico* permutation experiments with a bootstrapping significance threshold of 0.05. **(E-G)** Analyses of RNAi-mediated *zpg* knockdown LGs from late-second instar stage (N = 3), early-third instar stage (N = 4), and mid-third instar stage (N = 8). **(E)** Quantification of the number of spatially significant experiments in blood progenitors. See B. **(F)** Analysis of MEC magnitude. See C. **(G)** Number of validated hotspots per developmental stage. See D.

We previously showed that Ca^2+^ signaling appeared to evolve over larval development, correlating with the differentiation activity of progenitors (9). Specifically, we observed lower Ca^2+^ signaling propagation between neighboring cells and a reduced connectivity of the Ca^2+^ signaling network during early larval stages (9). To understand how Ca^2+^ signaling communities develop during hematopoiesis when progenitors show distinct proliferation and differentiation patterns (Fig. 3A), we expanded our analysis to the earlier stages of the late-second and early-third larval stages. Our analysis characterized a gradual build-up of signaling communities, in terms of both quantity and complexity, over the course of blood progenitor development. Specifically, both the fraction of experiments with significant local communities (Fig. 3B) and the magnitude of MEC (Fig. 3C) increased across the three stages in wild-type LGs. In contrast, RNAi-mediated *zpg* knockdown induced a decrease in both parameters of signaling communities (Fig. 3E-F), suggesting that the emergence of signaling communities was perturbed. Hotspots analysis showed a similar trend of gradual emergence of recurrent Ca^2+^ communities along the developmental trajectory with 0/4 statistically validated hotspots in the late-second, 3/6 in the early-third, and 8/14 in the mid-third stage(Fig. 3D). In contrast, no (0) hotspots were validated across all developmental stages of the Zpg-depleted LGs (Fig. 3G). Taken together, our data suggests that signaling communities and their recurrent activation (i.e., hotspots) emerge during, and evolve over, the course of larval development and that gap junctions are required for the developmental progression of these Ca^2+^ signaling communities in blood progenitors. This is also consistent with our previous observation showing that Zpg depletion increases blood cell differentiation (9), supporting a model where signaling communities coordinate blood progenitor behavior to maintain LG homeostasis during development.

## Discussion

There are numerous examples in the literature reporting synchronization and collective events in the context of cell signaling and behavior (13, 18, 19). A critical question that has remained underexplored is how does global, tissue-scale synchronization emerge from local cell-cell communication? More specifically, what are the intermediary steps involved in reaching the final synchronization state? In an attempt to provide some insight into the answers to these questions, we have previously described how endothelial monolayers synchronize Ca^2+^ signaling by gradually transitioning from local to global information spread (19). Other studies reported signaling waves propagating across long distances in a variety of systems and in the context of diverse functions (13, 20–22). However, these studies did not pinpoint a specific intermediate spatial scale between single-cell and collective signaling. Here, using *Drosophila* hematopoiesis as our model system, we were able to identify such an intermediate spatial scale. Our work elaborates on our previous findings that described the important role played by gap junctions in coordinating cellular signals in the LG (9). We now show that an intermediate spatial scale exists, involving transient gap junction-mediated Ca^2+^ signaling in the form of multicellular communities. Similar scale collective events were previously reported in the context of Erk signaling in epithelial cells, and Ca^2+^ signaling in the Madin-Darby canine kidney epithelium (13) suggesting that this could be a universal way to collectively organize the signaling activity of individual cells in a multicellular system.

A key feature in some of these transient communities was recurrent activation events that formed larger communication processing hubs that we call signaling hotspots. These hotspots had several important functional characteristics: 1) Their formation required the activity of Zpg-based gap junctions. 2) They acted as information hubs that were able to induce (i.e., transmit) and process (i.e., receive) collective signaling using mechanisms that operated both within (intrinsically) and outside (extrinsically) of the hotspot. 3) They exhibited repetitive interactions with their environment and were spatially heterogeneous. 4) There was an increased incidence of hotspots as the LG evolved and developed consistent with a role in the emergence of collective cell behavior. Each of these characteristics of the hotspots played an important functional role in shaping the signaling landscape within the LG. Overall, these findings reveal a novel mechanism whereby local cell-cell signaling propagation, through gap junctions, progresses into intermediate multicellular communities that integrate local information to achieve global population-wide synchronization during fly hematopoiesis.

Our observation that hotspots self-organize as information processing hubs in the blood progenitor population suggests that the hotspots perform a function that bears general resemblance to that performed by pacemaker cells, at the multicellular scale. A characteristic of pacemaker cells is their ability to coordinate the electrical or Ca^2+^ signaling activity of individual cells to guide collective decisions (18, 23, 24). Multicellular structures that are functionally and morphologically similar to pacemaker cells appear across diverse tissues, including Cajal interstitial cells in the gut (25), a sinoatrial node in the heart (24), and preBötC cells in the brain stem (26), indicating that it is a conserved module in living systems to regulate systemic homeostasis. We note three features of blood progenitor hotspots that resemble those found in pacemaking cells. First, as we previously proposed, blood progenitors form a small-world Ca^2+^ signaling network (9), where most cells are separated from each other by a small number of cell-to-cell transmission events thanks to a small subgroup of cells with high connectivity compared to other cells (12). Here, using ARCOS and *in silico* spatial permutation analysis, we directly demonstrated the existence of such hub-like network structures, or hotspots, within the blood progenitor population. Second, a well-known feature of pacemaker cells is their ability to integrate and segregate information between cells that are either external or internal to their signaling hub (27–29). Our study quantitatively illustrates that Ca^2+^ signaling in blood progenitors is organized into hotspots that are able to both receive and send information. Third, there are several functional analogies between the hotspots found in blood progenitors and cardiac pacemaker cells found in sinoatrial nodes. These include: (A) transfer of information across large distances and the ability to fine-tune the activities of a large group of cells (24), (B) highly synchronized multicellular activity that is often tied to function (9, 24, 30), (C) coordinated cell behavior that is dependent on gap junctions (9, 30), (D) self-organization and synchronization of local heterogeneous Ca^2+^ signals (24), and (E) intracellular Ca^2+^ signals in both systems are controlled by the same molecular machinery including gap junctions (9, 23, 30), SERCA pumps (9, 31), and ryanodine receptors (10, 31). These observations highlight similar design principles, both conceptual and functional, that allow LG blood progenitor hotspots and cardiac pacemaker cells to coordinate cells within a population.

Signaling hotspots highlight the spatial heterogeneity in intercellular Ca^2+^ information processing in the developing LG. How such heterogeneity develops in seemingly homogenous blood progenitors remains unknown. Heterogeneity in intercellular communication, even in the same cell population, can originate from intrinsic cell-to-cell variation in gene expression levels or protein modifications (32–34). Indeed, single-cell transcriptomic analysis on LGs showed that blood progenitors, which were previously considered as a homogenous population, exhibited a large variability in their gene expression profiles (35). The differences in their gene enrichment were used to classify progenitors into 6 main sub-clusters that showed distinct spatial distribution and gene expression profiles (35), suggesting that the difference in gene expression could contribute to the heterogeneity of Ca^2+^ signaling. Beyond gene expression or protein modifications, the positioning of the cells within the LG and in relation to other organs may lead to spatial heterogeneity between hotspots, by supporting different modes of cell-cell interaction (36). However, we were not able to identify a stereotypic spatial pattern in the hotspot location.

The emergence of hotspots from oscillating blood progenitors required a mechanism that coordinates their individual activities. Although we demonstrated that the function of Zpg-based gap junctions was indispensable in this process, the underlying mechanism remains unclear. We can envision several possible routes for the emergence of collective Ca^2+^ signaling hotspots in the blood progenitor population. According to theoretical, physics, and neural-based studies, routes giving rise to collective behaviors can be classified into four main categories (37): (A) Pacemaker cells, in this context cells that fire rhythmic signals, entraining other cells to oscillate or behave in a synchronized fashion (38). (B) Phase and/or frequency locking, where cells that naturally oscillate at different frequencies synchronize their behaviors by adjusting their phases and/or frequencies when coupled with other cells, a representative example being circadian neurons (39–42). (C) Oscillator death, where mathematical approaches and synthetic genetic clocks show that cells stop oscillating when coupled with other cells (37, 43). Therefore, decreasing the coupling strength permits the emergence of synchronized behavior. (D) Dynamic quorum sensing, where non-oscillatory cells start oscillating when a signaling molecule they secrete exceeds a critical concentration threshold in their environment, an example being yeast glycolytic oscillations (37). Comparing our data with the above four categories, we proposed that hotspot emergence likely involves a hybrid mechanism with both pacemaker-like and phase/frequency locking properties. First, we noticed that some progenitors were still able to produce Ca^2+^ spikes even in the presence of a high concentration of CBX (9), indicating that these cells spontaneously produce spikes without the need of neighbor connections. As discussed in the previous section (Hotspots act as information hubs), the progenitor hotspots show characteristics consistent with having pacemaker-like properties. Second, for the phase/frequency locking property, we found that the complexity and incidence rate of hotspots increased concomitant with animal development. This showed that hotspots are able to accommodate or incorporate new cells in a developing progenitor population. Our previous observations show that the number of gap junctions increased and the spiking frequency of blood progenitors was modulated during LG development (9). These two lines of evidence suggest that the newly incorporated cells, once coupled with other cells, changed their spiking frequency over time, consistent with the phase/frequency locking phenomenon. Overall, we suggested that the progenitor hotspots emerge by simultaneously utilizing the pacemaker-like and phase/frequency docking mechanisms. Taken together, our findings align with other recent studies that reported collective signaling in the spatial scale of multiple cells (13, 44, 45), suggesting a universal mechanism to collectively organize the signaling activity of individual cells in a multicellular system.

## Materials and Methods

### *Drosophila* genetics, stocks, and maintenance

All *Drosophila* stocks and crosses were maintained regularly on a standard cornmeal medium (recipe from the Bloomington *Drosophila* Stock Center) in vials or bottles at 25℃. The blood progenitor-specific Gal4 driver used was *Tep4-Gal4* (a kind gift from Dr. Lucas Waltzer, Université Clermont Auvergne, France). Other lines used were: *UAS-GCaMP6f* (RRID:BDSC_42747) and *UAS-zpg-RNAi* (RRID:BDSC_35607). Larvae were staged as follows: eggs were first collected 6∼8 hours after egg laying (AEL), late-second instar larvae were collected 68-72 hours AEL, early-third instar larvae were collected 72-80 hours AEL, and mid-third instar larvae (or wandering third instar larvae) were collected 96 hours AEL (9).

### Sample preparation and confocal imaging

To prepare live LG samples, larvae in desired stages were washed using Phosphate-Buffered Saline (PBS) three times (2 minutes each), quickly rinsed with 70% ethanol, washed again with PBS three times (2 minutes each), and dissected in the *Drosophila* Schneider’s medium (pre-warmed to room temperature 10 minutes prior dissection; ThermoFisher Scientific, 21720001). The dissected LG was mounted in the glass bottom dishes (MatTek Corporation, 35 mm, P35G-0-14-C, non-coated), covered with a 1% agar pad (Agar A, Bio Basic, FB0010, prepared in the Schneider’s medium), and stabilized with 1% agar spacers to prevent LG compression during live recordings (15). The dish was supplied with 2 ml Schneider’s medium over the agar pad for moisture and placed in a microscope incubator (TOKAI HIT, Catalog number: INU-ONICS F1) that maintains the temperature at 25°C during imaging. LG optical sections spaced by 1.5μm were imaged using a 40X oil immersion objective (numerical aperture 1.30, UPLFLN) on an Olympus inverted confocal microscope (FV1000) with a temporal resolution ranging from 2.3-6.7 seconds per frame (9).

To monitor real-time Ca^2+^ signals in blood progenitors, a genetically encoded Ca^2+^ sensor GCaMP6f (peak excitation ∼480 nm, peak emission ∼510 nm) was expressed. Fiji (46) was used to manually annotate circular ROIs around each progenitor cell according to the GCaMP6f activity. Raw GCaMP6f intensity values were extracted at the ROIs at individual time points (z-profile Fiji plugin) and exported to Excel (in .csv format). The obtained GCaMP6f signals of each cell were normalized, F’_t_ = (F_t_-F_min_)/(F_max_-F_min_) where F_t_ = raw GCaMP6f value at each time point, F_min_ and F_max_ = minimum and maximum GCaMP6f values of a cell, respectively) (9, ^15^). Time-lapse recordings were processed in Fiji and Fluoview (Olympus FV10-ASW 4.2) and the data was analyzed using Python. No stabilization or registration on images was performed. Intensities represented mean gray values.

To block gap junctions, live dissected LGs were incubated in 50 or 100 μM CBX (Sigma, CG4790) for 15 minutes, mounted in the Schneider’s medium with corresponding CBX concentration, and imaged immediately (9). For the CBX-washout experiment, LGs were incubated in 100 μM CBX for 15 minutes, rinsed in the Schneider’s medium twice (5 minutes each), mounted, and imaged immediately (9). A 1mM CBX stock was stored at −20 °C. Imaging settings were set identically across experiments.

### Transient communities detection and analysis

We applied ARCOS (13) to detect and quantify the Ca^2+^ collective signaling events in blood progenitors. We applied the ARCOS Python implementation (arcos4py, version 0.1.5) on the normalized time series for each inspected LG. We set neighborhoodSize to 14 μm, which represents about two cell diameters. minClsz, the minimum initial size for a cluster to be identified as a collective event, was set to 1. minTotalEventSize, the final size of the cluster at the end of the event, was set to 3 cells. This way we enforced a minimum cluster size of 3 cells while allowing asynchronous cell activations. nPrev, the maximal number of frames between different cell activations, was configured empirically to a maximum time lag of 15 seconds according to the temporal resolution. minDuration, the minimal time for a collective event, was set to 1 frame, enabling the detection of short-term co-occurring activations. Binarization parameters were set according to the default recommended values (13), with biasMet, smoothK and biasK set to “runmed”, 3 and 51, respectively. To minimize the detection of false activations, peakThr and binThr were empirically set to 0.3 and 0.4, respectively.

### Statistically validating local properties of collective signaling events

We designed a bootstrapping-based statistical test to reject the null hypothesis that the collective signaling events are non-local properties. This was achieved by repeating the following steps 1,000 times: (A) spatially shuffling the cells’ time series, which is equivalent to randomizing the cells’ locations; (B) applying ARCOS to the spatially shuffled time series; (C) recording the mean number of collective signaling events per cell (mean events per cell, MEC) across the spatially shuffled cells. The statistical significance was calculated as the fraction of spatially shuffled experiments where the MEC was equal to or exceeded the MEC of the observed (not shuffled) experiment. The MEC magnitude was calculated as the mean ratio between the experimentally observed MEC and the MEC of each of the spatially shuffled experiments, and indicates the MEC fold change in respect to excluding the spatial organization.

### Mean local cell density and mean cell activation

The *mean local cell density* was defined as the average number of cells within a square area of 14×14 µm^2^ surrounding each cell. The *mean activation rate* was defined as the average number of activations per cell per minute. Both measurements were calculated according to the mean value of all cells in each LG.

### Communities’ intercellular signaling propagation speed

The *intercellular signaling propagation speed* of a community was defined as the mean time difference between the activation of adjacent cells as a function of the distance between these cells (µm/second) in the context of the transient community. This community-specific measurement was pooled across all LGs within each experimental condition. To avoid confounding effects due to different temporal resolutions between experiments, we excluded experiments that had temporal resolution outside the range of 2.32-4 seconds per frame. This range maintains a sufficient and similar amount of LGs per treatment (*N_wild type late 2nd_* = 4; *N_wild type early 3rd_* = 5; *N_wild type mid 3rd_* = 4; *N_zpg RNAi late 2nd_* = 3; *N_zpg RNAi early 3rd_* = 3; *N_zpg RNAi mid 3rd_* = 3; *N_CBX_* _3.125_ = 2; *N_CBX_* _12.5_ = 1, *N_CBX washout_* = 4).

### Hotspots analysis

We defined LG_max_ as the maximal number of transient communities in which a single cell participated within a specific LG. We defined LG_threshold_ as the maximum between 5 and LG_max_, and marked all cells that participated in at least LG_threshold_ transient communities. For each connected component (in the neighborhood graph) group of adjacent cells above this threshold we calculated its convex hull and considered it as a *hotspot candidate*. To validate that a hotspot was not a result of random effects nor physical confounding factors (see Methods: Confounders analysis), we conducted a bootstrapping-based statistical test as follows. First, we matched at least 50% of the hotspot cells with other non-hotspot cells from the same LG, where each of the non-hotspot cells participated in at least the same number of transient communities as its matching hotspot cell. Second, we swapped the Ca^2+^ time series of each matched pair of hotspot and non-hotspot cells. Third, we employed ARCOS on the *in silico* spatially permuted LG to detect collective signaling events. Fourth, we recorded the MEC for the permuted hotspot cells. Fifth, we repeated these four steps for each hotspot up to 1000 times, hotspot candidates with at least 100 different *in silico* spatially permuted LGs were considered for the bootstrapping-based significance test. For each hotspot candidate, the statistical significance was determined as the percentage of *in silico* permutations that yielded equal or greater MEC values compared to the original non-permuted LG. A hotspot candidate with a p-value ≤ 0.05 was considered as a *validated hotspot*.

### Interactions between hotspots and their surrounding environment

We quantified the interaction between cells within hotspots and their adjacent non-hotspot cells, and measured the temporal ordering of the cells’ activation. *Hotspot community* was each transient community that included at least one hotspot cell. For each hotspot, we calculated the ratio between the number of hotspot communities involving both hotspot and non-hotspot cells to the total number of hotspot communities (also including hotspot-exclusive cells). This ratio represents the probability of hotspot cells interacting, via a transient community, with non-hotspot cells.

The direction of interaction between hotspot and non-hotspot cells was defined as whether a hotspot community was initiated by a hotspot or a non-hotspot cell. This analysis focused on hotspot communities involving at least one non-hotspot cell. We defined two measurements for directionality: (A) The fraction of hotspot communities that were initiated by hotspot cells. For this measurement, we excluded hotspot communities that were initiated by both hotspot and non-hotspot cells that appeared in the same time frame, because of the ambiguity to which cell initiated the community. (B) For each hotspot transient community, we considered all cell pairs comprising one hotspot cell and one non-hotspot cell, within a distance ≤ 14 µm from one another. We calculated the *transmission probability* as the fraction of such pairs where the hotspot cell was activated before the non-hotspot cell.

The hotspot size was defined as the number of cells participating in the hotspot. The proportion of hotspot cells in transient communities was defined as the fraction of hotspot cells in a community. This proportion was averaged across all hotspot communities to define the average proportion of hotspot cells in transient communities, which was used as the expected probability of a hotspot cell to be the initiator of a hotspot transient community, under the assumption of random activation order of cells within a community.

### Statistical analysis

Pearson correlation (scipy.stats.pearsonr) was used to measure the correlation between the Ca^2+^ signals of blood progenitors (see Fig. 1B-C) and the correlation between MEC rate, mean local cell density, and mean cell activation rate (see Fig. S3, Fig. S4). Bootstrapping was applied in the spatial shuffle analysis (e.g., Fig. 1E) and the hotspot shuffle analysis (e.g., Fig. 2C). Fisher’s exact test (scipy.stats.fisher_exact) was used to measure the differences between different experimental conditions (treatments) in terms of the amount of spatially significant LGs (e.g., Fig. 1F-G, Fig. 3B, Fig. 3E). Fisher’s exact test was chosen due to the small sample size in each experimental condition, and due to the categorical nature of the data. Kruskal-Wallis test (scipy.stats.kruskal) was used to measure the difference between the distributions of cell pair Pearson correlation of Ca^2+^ signals (Fig. 1C, Fig. S1B), magnitude of MEC (Fig. 1H, Fig. 3C, Fig. 3F), community-level information spread rate (Fig. 1I), and distance distribution comparison (Fig. S4D) across experimental conditions. Non-parametric Kruskal-Wallis test was chosen due to the varying sample sizes across different experimental conditions and due to the unknown underlying distribution of our data. All significance tests were carried out with an α-value of 0.05, considering * - p < 0.05, ** - p < 0.01, *** - p < 0.001, **** - p < 0.0001.

## Software and data availability

We are currently organizing our source code and will make it publicly available as soon as possible (before journal publication).

## Funding and Acknowledgments

The authors acknowledge the Bloomington *Drosophila* Stock Center. The authors thank Dr. Lucas Waltzer for providing fly stocks. Research at the Zaritsky laboratory is supported by the Israel Council for Higher Education (CHE) via the Data Science Research Center, Ben-GurionUniversity of the Negev, Israel, by the Israel Science Foundation (ISF, grantNo. 2516/21), by the Israel Ministry of Science and Technology (MOST), by the Wellcome Leap Delta Tissue program, by the German-Israeli Foun-dation (GIF)-Nexus, and by the Rosetrees Trust. Research in the lab of G.T. is supported by the Canadian Institutes of Health Research (Project Grant PJT-156277). K.Y.L.H is supported by a 4-Year Doctoral Fellowship from the University of British Columbia. The authors thank Dr. Bo Sun, Dr. Dagan Segal, and Dr. Julia Mack for critically reading the manuscript.

## Author Contribution

AZ, SBD, and GT conceived the study. KYLH designed the experimental assay and performed all experiments. SBD developed analytic tools, analyzed, and interpreted the data with the help of KYLH. AZ and GT mentored SBD and KYLH. All authors wrote and edited the manuscript and approved its content.

