## Supplemental figures and figure legends Figure for "Formation of recurring transient Ca^2+^-based intercellular communities during *Drosophila* hematopoiesis"

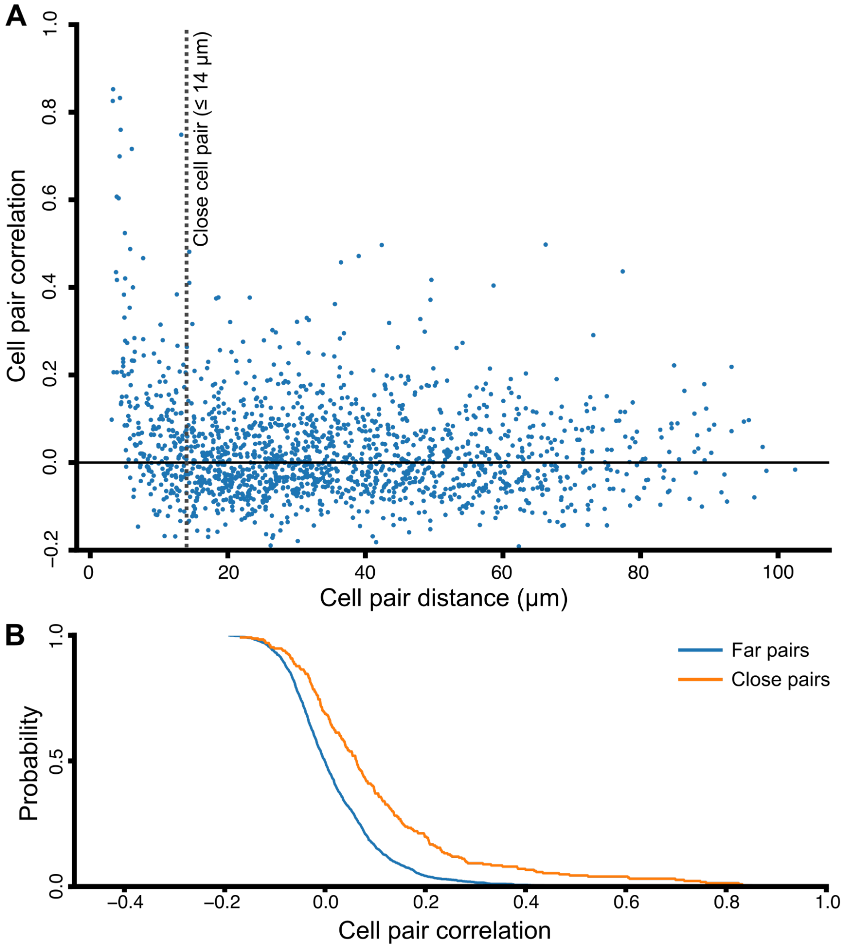


**Figure S1. Close blood progenitor pairs are more synchronized in their Ca2+ activities than distant cell pairs. (A)** Spatial analysis of all blood progenitor pairs in their temporal $Ca^{2+}$ pattern. The temporal correlation of $Ca^{2+}$ signals between cell pairs was correlated with the distance (in μm) between cell pairs. Each blue dot represents a cell pair. $N_{cells}$= 57, $N_{pairs}$= 1596, Pearson correlation between the $Ca^{2+}$ correlation of cell pairs and their corresponding distance = -0.15, p-value < 0.0001. **(B)** Cumulative distribution of Pearson correlation of the close (orange; N=227, μ=0.101, σ=0.181) and far (blue; N=1369, μ=0.016, σ=0.097 ) blood progenitor pairs (same pairs as in A). Each value $F_g(x)$ in the plot is the probability of a pair in group $g$ having a Pearson correlation coefficient greater than $x$. Kruskal-Wallis statistical test verified a significant difference between the two distributions (p-value < 0.0001).

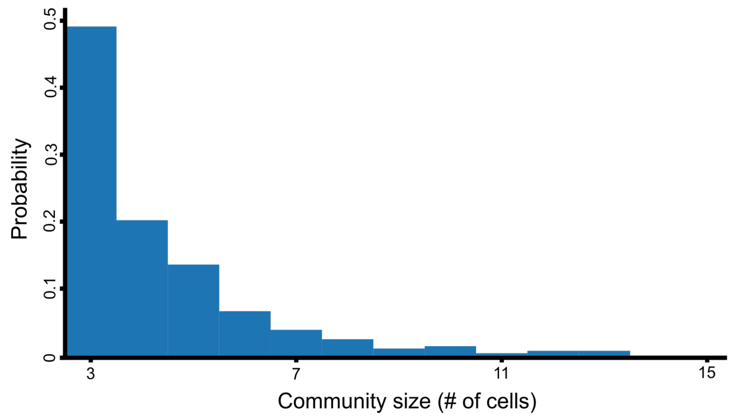


**Figure S2. Distribution of community sizes.** Transient communities pooled across 12 wild-type LGs. Each bar is the probability of a community involving the corresponding number of cells. $N_{communities} = 288$, $\mu_{size} = 4.302$.

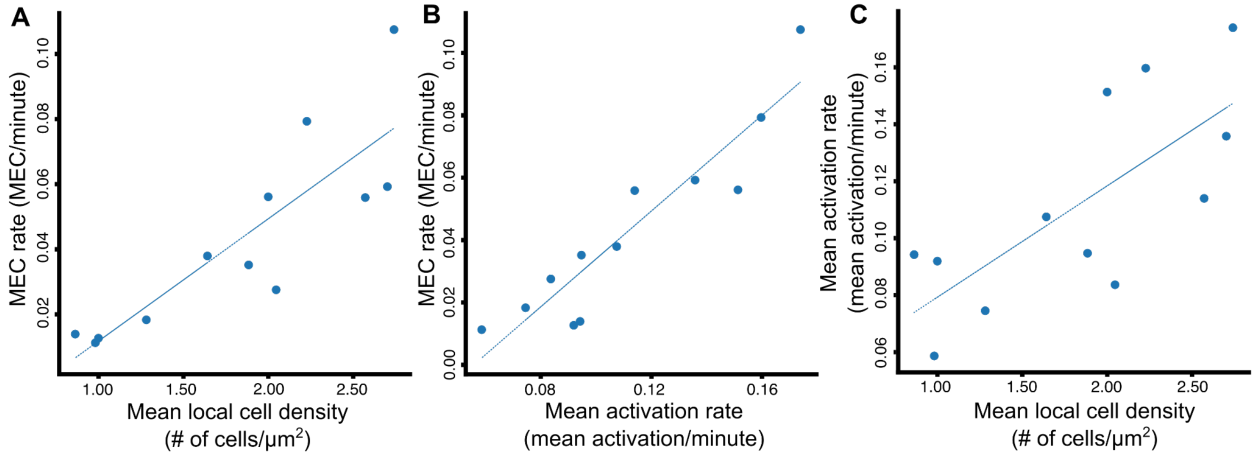


**Figure S3**. **Correlation between potential confounding factors**. Each data point reflects the mean value of the corresponding measurements across all cells in an experiment, N = 12 wild-type LGs. MEC rate is the number of MECs per minute. The mean activation rate is the average number of activations per cell per minute. Mean local cell density is the average number of cells within an area of 14x14 $\mu m^2$ surrounding each cell. The line represents linear fit. **(A)** Pearson correlation (coefficient = 0.854, p-value = 0.0004) between MEC rate and the mean local cell density **(B)** Pearson correlation (coefficient = 0.929, p-value < 0.0001) between MEC rate and mean activation rate. **(C)** Pearson correlation (coefficient = 0.735, p-value = 0.0065) analysis between mean activation rate and mean local cell density.

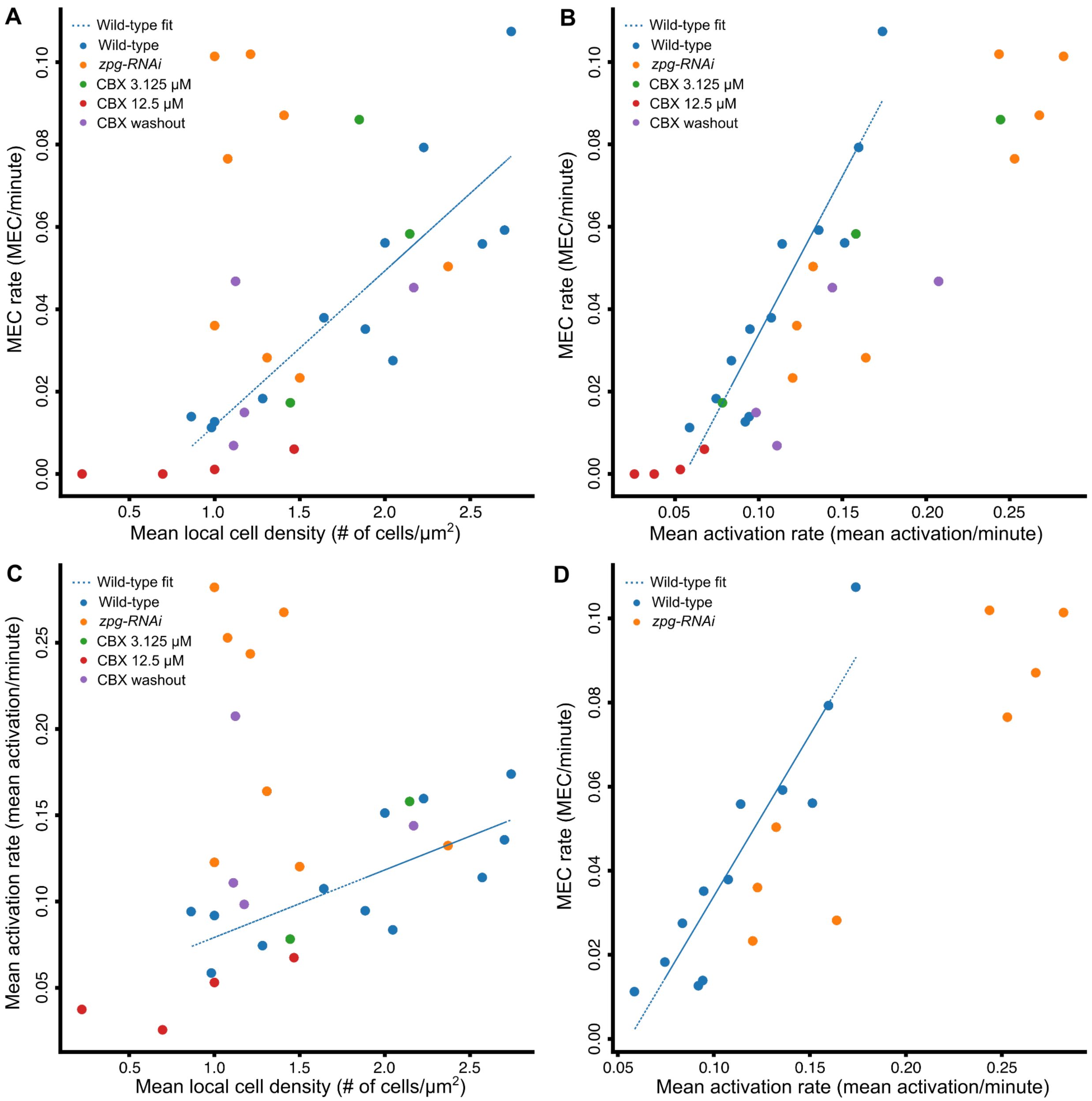


**Figure S4**. **Correlation between potential confounding factors in gap junction inhibition experiments**. Each data point reflects the mean value of the corresponding measurements across all cells in an experiment, with wild-type LGs in blue (N=12), RNAi-mediated knockdown of *zpg* in orange (N=8), 3.125 µM CBX in green (N=3), 12.5 µM CBX in red (N=4), and CBX washout in purple (N=4). The blue line represents the linear fit based on the wild-type LGs. **(A)** Pearson correlation (coefficient = 0.487, p-value = 0.0055) between MEC rate and the mean local cell density. (**B**) Pearson correlation (coefficient = 0.876, p-value < 0.0001) between MEC rate and mean activation rate. **(C)** Pearson correlation (*zpg RNAi* data was excluded from the calculation of the Pearson correlation, see panel D; coefficient = 0.573, p-value = 0.0042) between mean activation rate and mean local cell density. **(D)** MEC rate and mean activation rate for wild-type (blue; N=12) and RNAi-mediated knockdown of *zpg*

(orange; N=8) LGs. Data as in panel B. The distance of each *zpg RNAi* LG from the wild-type linear fit was calculated as the subtraction between the observed and the corresponding linear-fit MEC rate ($\mu_{wild\ type}$ = (- 9.83$e$)$^{-18}$, $\sigma_{wild\ type}$ = 0.01, $\mu_{RNAi\ zpg}$ = - 0.046, $\sigma_{RNAi\ zpg}$ = 0.025). Kruskal-Wallis statistical test verified a significant difference between the distance distributions of the two experimental groups (p-value = 0.0009).

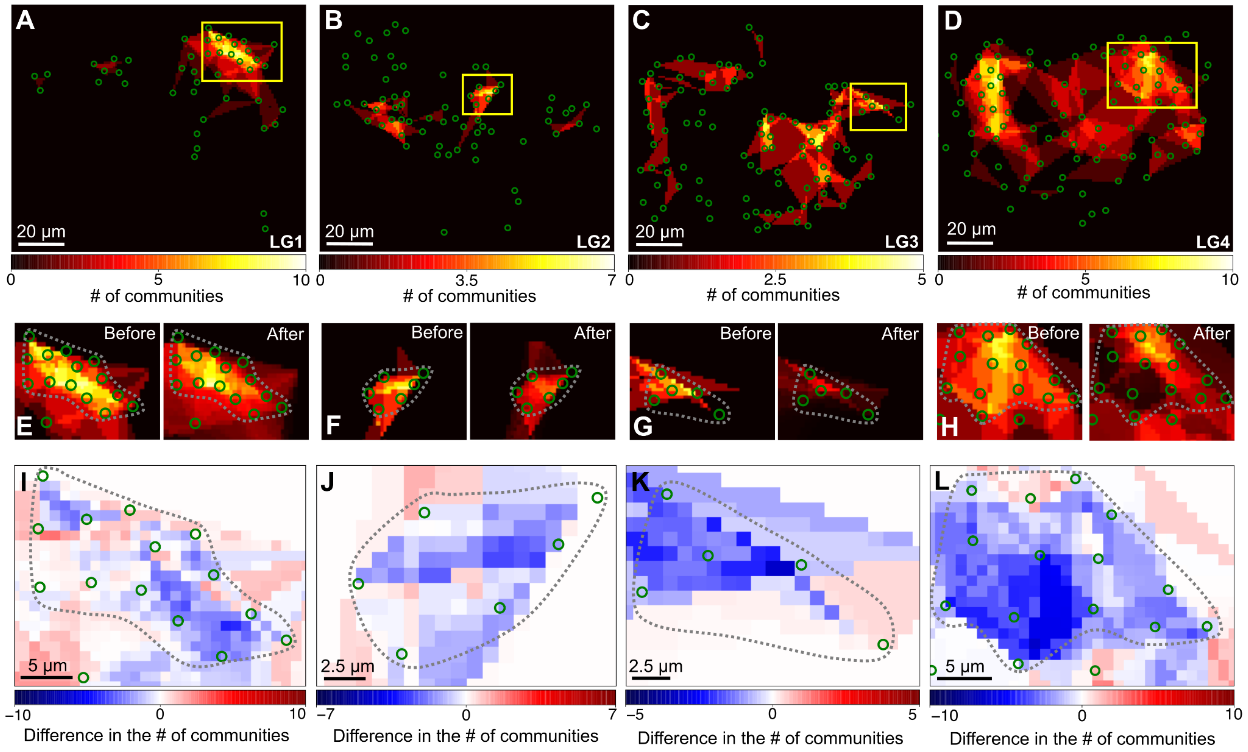


**Figure S5. Spatial *in silico* permutations of hotspot cells reduce communication in a hotspot. (A-D)** Visualization of the integrated number of transient communities each cell participated in over time in different wild-type LGs. Green circles: the center of each blood progenitor. Color-coded legend: the number of communities. Yellow regions of interest mark hotspots. **(E-H)** Hotspots (dashed lines) before and after *in silico* permutation. Each hotspot corresponds to the hotspot above it in panels A-D **(I-L)** Signed change in the number of transient communities within a hotspot's (dashed line) cells with respect to the mean number following *in silico* permutations. Each hotspot corresponds to the hotspots above it in panels A-D and E-H.

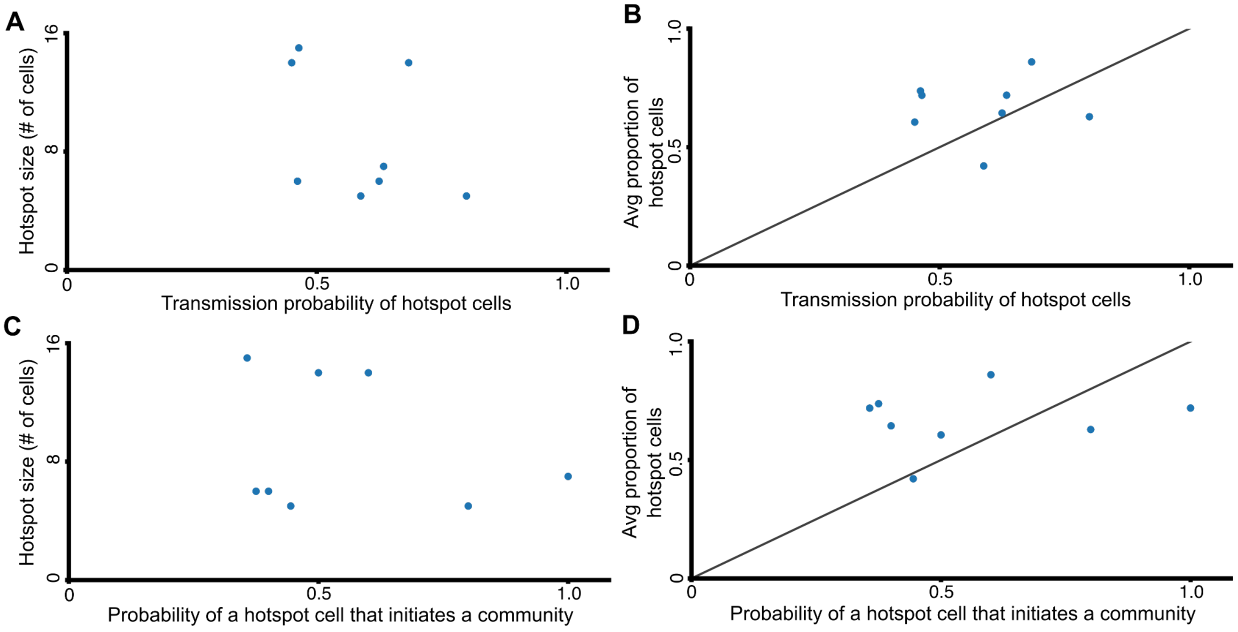


**Figure S6. Interactions of hotspots with their surrounding cells did not have a predominant direction.** Each data point reflects the probability across all transient communities in a statistically significant validated hotspot. N = 8 hotspots pooled across wild-type LGs. The average proportion of hotspot cells in transient communities (y-axis in B and D) was defined as the mean of the ratio between the number of hotspot cells in a community and the total number of cells in that community. The diagonal (y = x) indicates the situation where the average proportion of hotspot cells corresponds to their probability of transmitting a signal, i.e., activating before non-hotspot cells in the same transient community (B), or initiating a transient community, i.e., activated first in the transient community (D). Transmission probability was calculated from all pairs of adjacent hotspot and non-hotspot cells in a common transient community (see Methods). **(A-B)** Probability of signal transmission from hotspot cells to non-hotspot cells as a function of hotspot size (**A**) and as a function of the average proportion of hotspot cells in transient communities (**B**). **(C-D)** The probability of a hotspot cell to initiate a transient community as a function of hotspot size (**C**) and as a function of the average proportion of hotspot cells in transient communities **(D)**.

### Supplemental video legends

**Video S1:** $Ca^{2+}$ signaling dynamics in blood progenitors of a wild-type LG, visualized using GCaMP6f (green). Timestamp is measured in seconds.

**Video S2:** $Ca^{2+}$ signaling dynamics of a wild-type LG, highlighting three transient communities (each community is annotated by a red, white, or yellow polygon). Timestamp is measured in seconds.

**Video S3:** The formation and disintegration of the 6-cell transient community shown in Fig. 1D. Timestamp is measured in seconds.
